# Long distance retrograde degeneration of the retino-geniculo-cortical pathway in homonymous hemianopia

**DOI:** 10.1101/750208

**Authors:** Alekya P. Rajanala, Mohammad A. Shariati, Yaping Joyce Liao

**Author notes:** Corresponding author (YJL).

## Abstract

Long-distance retrograde degeneration of the retino-geniculo-cortical pathway has been described in humans and animal models following injury to the brain. In this study, we used optical coherence tomography (OCT) to measure the severity and timing of retrograde degeneration after post-chiasmal visual pathway lesions in patients with homonymous hemianopia. We performed a retrospective study of 69 patients with homonymous hemianopia and analyzed high quality OCT macular ganglion cell complex (GCC) and retinal nerve fiber layer (RNFL). Patients with lesions involving the optic tract and thalamus were included in the anterior group, while patients with lesions of the occipital lobe were included in the posterior group. Statistical significance was determined using Mann-Whitney U test and Wilcoxon test. We found that in patients with homonymous hemianopia, those with anterior lesion exhibited earlier and more severe thinning compared with the posterior group. In fact, thinning can occur within 2 months after insult in the anterior group. Within 6 months of onset, the anterior group exhibited about 5 times more hemi-macular GCC thinning than those with acquired lesions of the posterior visual pathway (P = 0.0023). Although the severity of hemi-macular GCC thinning was different, the majority of hemi-macular thinning occurred within the first 6 months in both groups. Beyond 2 years, thinning in those with acquired anterior and posterior lesions was minimal, except in a small number of patients with multiple insults to the occipital lobe. In conclusion, using OCT, we measured the severity and rate of long-distance, retrograde degeneration in patients with homonymous hemianopia. Homonymous hemi-macular thinning after optic tract and thalamic injury was more severe and occurred earlier compared with thinning after occipital lobe insult via trans-synaptic degeneration. The presence of severe hemi-macular degeneration on OCT provides objective evidence that localizes the lesion to the post-chiasmal anterior visual pathway.

## Introduction

The retino-geniculo-cortical visual pathway is a 3-neuron long-distance white matter pathway, and insult to any part of this pathway leads to devastating vision loss. Although vision loss in one eye can be compensated somewhat by using the remaining good eye to function, injury posterior to the optic chiasm is associated with homonymous hemianopia, which means one lesion in the brain leads to loss of the left or right side of vertical midline in both eyes. Homonymous visual field defect is most commonly due to stroke, and of these, most are due to occipital stroke [1–3]. Among 850 patients with 904 events causing homonymous hemianopia that were confirmed by neuroimaging, 70% were due to stroke (4% bilateral), and 30% were due to other conditions, such as trauma (14%), brain tumor (12%), neurosurgical procedures (2.5%), and multiple sclerosis (1.5%) [4]. Homonymous hemianopia cannot be easily compensated since both eyes are affected, and it leads to severe disability because patients can lose their driving eligibility. There is also disability due to hemianopic dyslexia [5, 6], deterioration of mental health [7, 8], and impairment of activities of daily living [9–11].

Retrograde long-distance degeneration occurs after visual field loss. This has been described after optic neuritis [12] and chiasmal compression [13], which involves degeneration of the optic nerve (up to about 5 centimeters) and loss of the retinal ganglion cell somata. After occipital lobe insult, degeneration across the retino-geniculo-cortical pathway can occur in a retrograde fashion trans-synaptically over 20 centimeters [14, 15]. The most definitive studies of retrograde trans-synaptic degeneration have been done with ablation of the striate cortex in non-human primates, which have shown that within 14-100 days, retrograde degeneration extends from the occipital lobe into the lateral geniculate, optic tract, and the eye [16, 17]. Retrograde trans-synaptic degeneration of the visual pathway has also been described in human autopsy and MRI studies [18–21].

Advances in optical coherence tomography (OCT) means that we now have a reliable *in vivo* method of measuring long-distance, retrograde degeneration of the visual pathway in humans at point-of-care locations such as the eye clinic [22, 23]. Commercial spectral-domain OCT machines provide resolution of 3-5 microns, which allow for quantification of individual retinal layers comparable to that of retinal histology [24, 25]. Advances in OCT segmentation means we can routinely segment the thickness of the layer containing retinal ganglion cells (ganglion cell layer or GCC) and the thickness of the layer containing unmyelinated retinal ganglion cell axons (retinal nerve fiber layer or RNFL). Modern OCT machines also have eye tracking, which allows for good inter-scan reproducibility [26–28], which facilitates comparison of OCT measurements over time. OCT measurements have been useful in measuring changes in thickness in different retinal layers due to optic nerve diseases, such as glaucoma, optic neuritis, ischemic optic neuropathy, papilledema, and traumatic optic neuropathy [29–32].

In this study, we aimed to measure the severity and rate of retrograde degeneration after onset of homonymous hemianopia. We recruited patients with homonymous visual field defect and used OCT to measure the severity and timing of retrograde degeneration after injury to the post-chiasmal visual pathway. In particular, we compared measurements between lesions affecting the anterior post-chiasmal pathway, involving the optic tract and the thalamus, with lesions of the posterior post-chiasmal pathway, involving the occipital lobe. Understanding how an insult to one region of the brain affects white matter tracts and neuronal survival elsewhere is important from a biological as well as therapeutic standpoint.

## Methods

### Patient Selection

Our study was approved by the Stanford Institutional Review Board. We performed a retrospective case review of 79 consecutive patients seen in 2012-2017 and identified 69 patients (51% male), 138 eyes, and 73 events with homonymous visual field defect. Thirty-one patients (45%) had left-sided lesions, and 34 patients (49%) had right-sided lesions. Four patients (6%) had with bilateral, sequential strokes. Thirty-six patients (52%) had 2 or more OCTs. Mean age of all patients was 54.7 ± 2.5 years (median 57 years, range 15-91 years). Mean age at onset of disease was 41 ± 4.7 years for the anterior group and 61.3 ± 2.5 years for the posterior group. Mean age for the congenital group was 41.4 ± 8.6 years. All patients were followed for at least one year, and 21 (30%) were followed for 2 years or more.

We included patients who had formal visual field testing to confirm presence of homonymous visual field defect in both eyes. This was done with automated static perimetry (Humphrey Visual Field Analyzer, Carl Zeiss Meditec, Germany) or manual perimetry (Goldmann Visual Field model, Haag-Streit, USA). Each patient had to have average mean deviation ≤ -3 dB on static perimetry or at least partial quadrantanopia on Goldmann visual field testing. Those with visual field loss affecting less than one quadrant (e.g. homonymous scotomas) were excluded. We excluded patients with ophthalmic or neurological conditions of the visual pathway that could potentially affect the measurements.

Brain imaging was performed in all patients to confirm location and etiology of the visual field loss. Lesions involving the anterior post-chiasmal visual pathway, which involved the optic tract and the lateral geniculate nucleus, were considered to be in the anterior group. Lesions not involving the latter structures and primarily involving the occipital lobe were in the posterior group. One patient who had both anterior and posterior involvement due to multi-focal stroke was classified into the anterior group because of the involvement of the anterior structures. All patients with posterior cerebral artery strokes were included in the posterior group, regardless of stroke size. Four patients (6%) had bilateral posterior lesions and therefore did not have an “unaffected” side; these patients’ OCT and visual field data were analyzed without assigned control values.

We analyzed Humphrey Visual Field (HVF) data by mean deviation (dB) of the nasal and temporal hemifield using the Hood-Kardon linear model [22, 23]. The contralateral temporal field and ipsilateral nasal field were averaged as “abnormal” and the contralateral nasal field and ipsilateral temporal field were averaged as “control”. There was no difference in visual field severity between the anterior and the posterior group (anterior: -8.1 ± 2.2 dB, N = 7; posterior: -11.6 ± 1.6 dB, N = 33; P = 0.4011, Mann-Whitney).

### Optical Coherence Tomography Data Acquisition

To assess changes in retinal thickness over time, we performed spectral-domain optical coherence tomography (OCT) (Cirrus HD-OCT model, Carl Zeiss Meditec, Dublin, CA, USA). Because age-related OCT thinning is relatively modest, at 1 µm of GCC or 2 µm of RNFL every decade [24, 25], the changes we measure over time likely reflected changes corresponding to the homonymous visual field defect. We performed the Macular Cube 512 x 128 and the Optic Disc 200 x 200 scans per manufacturer’s instructions. We used high-resolution optical coherence tomography with eye-tracking capabilities and followed patients longitudinally as early as 1 week and up to 5 years after lesion onset. Only images with signal strength of 7 or above were included in the analysis. A total of 225 OCT (112 GCC; 113 RNFL) scans were performed. Nineteen scans were rejected based on poor segmentation, and 102 GCC and 104 RNFL scans were included in our analyses. All calculations were automatically segmented using the Zeiss algorithm and visually inspected to ensure appropriate segmentation.

### Macular Ganglion Cell Complex OCT Analysis

On OCT, macular GCC thickness was measured as the combined thickness of the ganglion cell layer and the inner plexiform layer. We calculated the GCC thickness corresponding to the side of visual field loss (the “abnormal” side) and the side with normal visual field (the “control” side) by averaging the measurements in the two eyes [33, 34]. For example, in a patient with left homonymous visual field defect, the abnormal side was calculated as the average of the right eye superior temporal and inferior temporal hemi-macular GCC and the left eye superior nasal and inferior nasal GCC. The control side was the average of the right eye superior nasal and inferior nasal GCC and the left eye superior temporal and inferior temporal GCC. To calculate the amount of GCC thinning for each subject, we subtracted the abnormal side from the control side [33]. To compare GCC thickness in the abnormal and control side in patients in the anterior and posterior groups, only 1 measurement per subject was used. If a patient had more than one OCT measurements, then the most recent OCT measurement was used. The range of abnormal GCC thickness was 43.5 to 74.5 µm in the anterior group, 45.8 to 91.3 µm in the acquired posterior group, and 43 to 77.5 µm in the congenital/incidental posterior group. Four patients in the anterior group, 4 patients in the posterior acquired group, and 1 patient in the posterior congenital group had very thin measurements that we visually confirmed was correctly measured.

To calculate the crossed GCC thickness measurements, we calculated the average thickness of the superior nasal and inferior nasal GCC. The non-crossed GCC thickness was calculated as the average thickness of the superior temporal and inferior temporal GCC. For example, in left homonymous visual field defect patients, the abnormal crossed GCC thickness was the average superior nasal and inferior nasal GCC in the left eye, and the control crossed GCC thickness was the average superior nasal and inferior nasal GCC in the right eye. In the same patient subgroup, the abnormal non-crossed GCC thickness was the average superior temporal and inferior temporal GCC in the right eye, and the control non-crossed GCC thickness was the average superior temporal and inferior temporal GCC in the left eye.

### Retinal Nerve Fiber Layer OCT Analysis

RNFL thickness measurement was calculated as the average of the crossed and non-crossed fibers in both eyes [33]. Unlike the GCC, which demonstrates clear segregation of the crossed and the non-crossed visual pathways into the nasal and temporal retinae, respectively, all 4 quadrants of the RNFL contain a combination of crossed and non-crossed fibers. However, the nasal quadrant is known to contain preferentially more crossed fibers, while the superior, inferior, and temporal quadrants contain a majority of non-crossed fibers [23, 35]. Therefore, we analyzed crossed fibers using nasal quadrants, with nasal retina contralateral to the brain lesion as “abnormal” and ipsilateral nasal retina as “control.” We then analyzed non-crossed fibers as an average of superior and inferior quadrants of RNFL; superior and inferior retina ipsilateral to the brain lesion were defined as “abnormal” and contralateral superior and inferior retina as “control”. The temporal quadrant was not included in this particular analysis because it is relatively much thinner than the superior and the inferior quadrants. The non-crossed fibers were then separately analyzed again as an average of superior, inferior, and temporal quadrants of RNFL; ipsilateral superior, inferior, and temporal retina were defined as “abnormal” and contralateral superior, inferior, and temporal retina as “control” [33].

### Statistical Analysis

We used the non-parametric Mann-Whitney U test and the non-parametric Wilcoxon matched-pairs signed rank test in Prism (GraphPad; La Jolla, CA), and used Prism to plot the data. The cut-off for significant values was set at P < 0.05. All data were presented as mean ± standard error of mean (SEM).

## Results

### Patient Characteristics

We performed a case-control study of 69 patients with homonymous hemianopia in order determine how optical coherence tomography (OCT) measurements change over time (Table 1). There were 15 patients with lesions involving the anterior post-chiasmal visual pathway, including the optic tract and thalamus (lateral geniculate nucleus). This group was called the “anterior” group because of involvement of the immediate post-chiasmal structures, even if the insult included nearby structures (e.g. temporal lobe). All lesions in the anterior group were acquired, and the most common causes included traumatic brain injury, tumors, and hemorrhage. There were 54 patients with lesions involving the posterior post-chiasmal visual pathway, including the occipital lobe and sometimes occipito-temporal or occipito-parietal lobes (tumors, arteriovenous malformation). This group was called the “posterior” group because the lesions involved structures beyond the lateral geniculate nucleus and did not involve the immediate post-chiasmal structures. Forty-seven patients in the posterior group had acquired lesions, most commonly due to posterior cerebral artery strokes and tumors. Seven patients in the posterior group had congenital or incidentally found occipital atrophy and were analyzed separately.

**Table 1.**
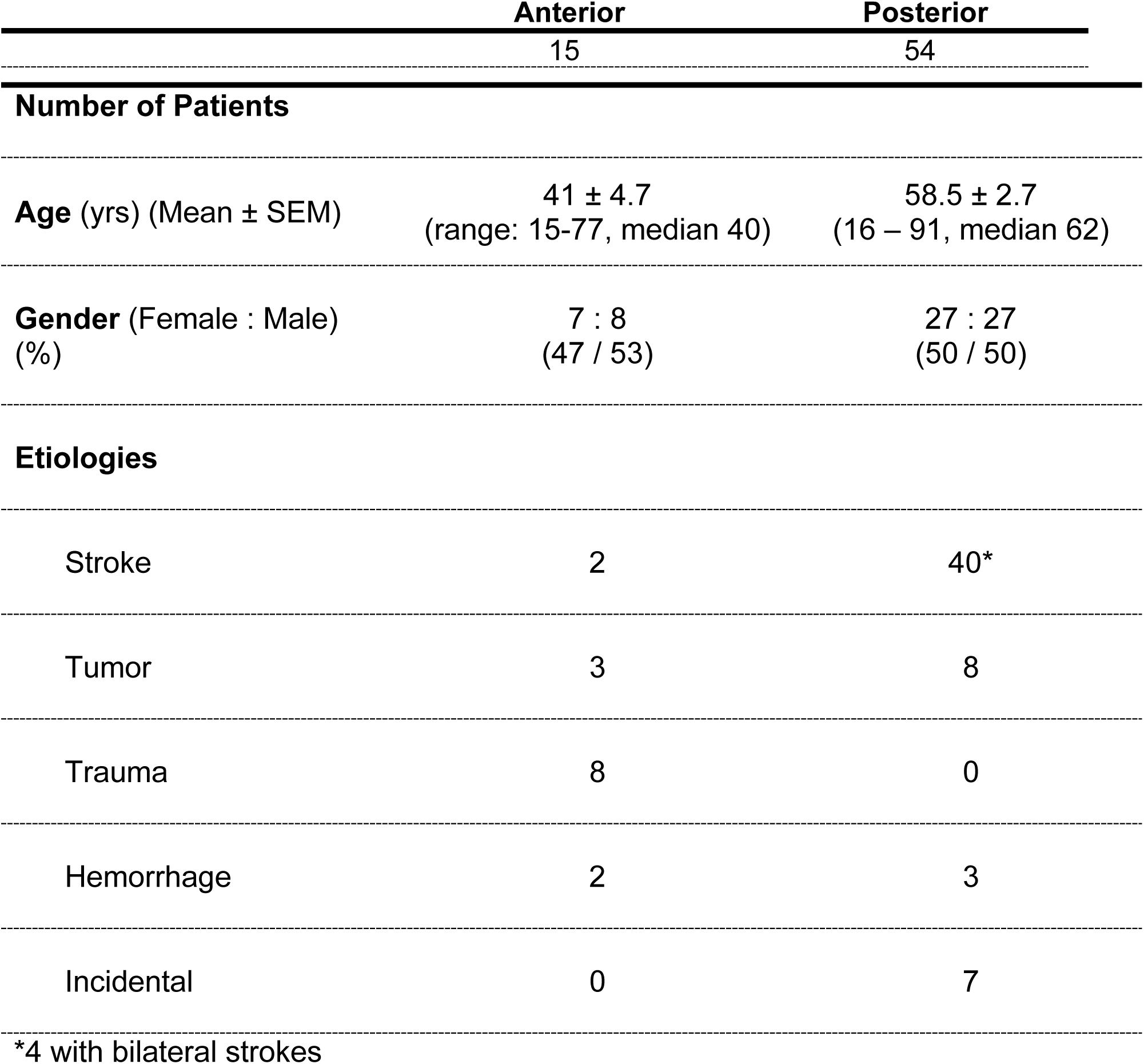
Clinical Features of Patients with Homonymous Visual Field Defect

| Anterior | Posterior |
| --- | --- |
| 15 | 54 |

Number of Patients
|  |  |  |
| --- | --- | --- |
| **Age** (yrs) (Mean ± SEM) | 41 ± 4.7<br>(range: 15-77, median 40) | 58.5 ± 2.7<br>(16 – 91, median 62) |
| **Gender** (Female : Male)<br>(%) | 7 : 8<br>(47 / 53) | 27 : 27<br>(50 / 50) |

Etiologies
|  |  |  |
| --- | --- | --- |
| Stroke | 2 | 40* |
| Tumor | 3 | 8 |
| Trauma | 8 | 0 |
| Hemorrhage | 2 | 3 |
| Incidental | 0 | 7 |

\*4 with bilateral strokes

An example patient in the anterior group is a 51-year-old man with acute onset vision loss due to hypertensive thalamic hemorrhage (Fig 1A, 1C, and 1E). His systolic blood pressures were over 200 mm Hg, and the hemorrhage involved the right optic tract in the thalamus and lateral geniculate nucleus as seen on brain magnetic resonance imaging (MRI) (Fig 1A). Kinetic perimetry with Goldmann visual field test revealed homonymous left inferior quadrantanopia (Fig 1C). OCT macular GCC analysis 2 months after onset revealed prominent thinning of the corresponding macula in a homonymous, hemi-macular pattern, which involved the temporal macula in the right eye and nasal macula in the left eye (Fig 1E).

**Fig 1:**
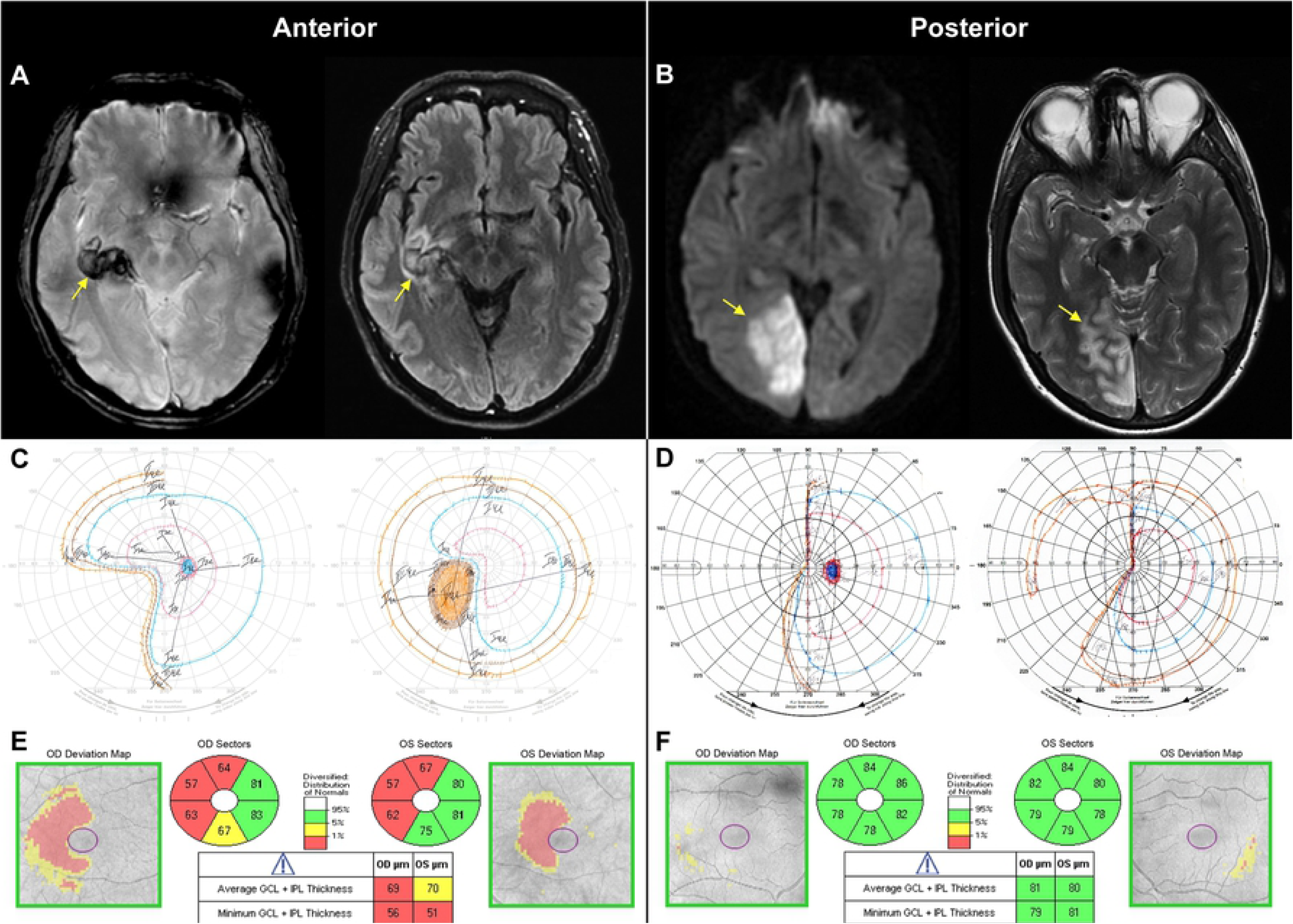
Representative example of homonymous visual field defect after insult involving the anterior (A, C, E) or the posterior (B, D, F) post-chiasmal visual pathway. **A.** Brain magnetic resonance imaging (MRI) axial gradient recall echo (GRE) and axial FLAIR (fluid-attenuated inversion recovery) scans showing right thalamic hemorrhage associated with hypertension (yellow arrows). The hemorrhage and edema involved the right optic tract within the basal ganglia and the lateral geniculate nucleus. **B.** Brain MRI axial diffusion weighted imaging (DWI) and axial T_2_-weighted scans showing stroke (yellow arrows) in the right posterior cerebral artery territory. **C**. Kinetic perimetry for anterior patient showing homonymous left inferior quadrantanopia. **D.** Kinetic perimetry for posterior patient showing homonymous left hemianopia. **E**. OCT macular GCC analysis for the anterior patient 2 months after onset of vision loss exhibited homonymous hemi-macular thinning involving the right eye temporal and left eye nasal maculae. There was a GCC thickness difference of 21.5 µm between the abnormal side (average of right eye temporal GCC and left eye nasal GCC) and control side (average of right eye nasal GCC and left eye temporal GCC) (abnormal: 59.8 µm, control: 81.3 µm). **F.** OCT macular GCC analysis for the posterior patient 3 months after onset of vision loss exhibited no thinning in either eye.

An example patient in the “posterior” group is a 38-year-old woman with systemic lupus erythematosus who developed a right posterior cerebral artery stroke in the setting of 5 days of fever, vomiting, and diarrhea after a dental procedure (Fig 1B, 1D, and 1F). Brain MRI revealed restricted diffusion and hyperintensity on T2-weighted images (Fig 1B). Her Goldmann visual field test showed a left homonymous hemianopia (Fig 1D). In contrast to the anterior patient, her OCT macular GCC performed 3 months after onset revealed no thinning in either eye (Fig 1F). Thus, there was a dramatic difference between the anterior patient, who exhibited severe macular GCC thinning, and the posterior patient, who exhibited normal macular GCC measurements within 2 months after insult.

### Significantly greater GCC thinning in anterior group

To determine whether the difference in macular GCC was present in all patients in the anterior and the posterior groups, we included one mean macular GCC measurement per patient and compared GCC thickness corresponding to the side of visual field loss (“abnormal” side) and the side corresponding with normal visual field (“control” side) in patients with acquired lesions (see Methods). In the anterior group, the abnormal side had 23.1 µm thinner GCC relative to the control side (abnormal: 56.5 ± 2.6 µm, control: 79.6 ± 1.5 µm, N = 15, P < 0.0001, Wilcoxon signed-rank test) (Fig 2B), while the posterior group, had 6.6 µm thinner GCC in the abnormal side (abnormal: 69.9 ± 1.6 µm, control: 76.5 ± 1.2 µm, N = 41, P < 0.0001, Wilcoxon). This means that among patients with acquired lesions, the anterior group had 3.5 times greater GCC thinning between the abnormal and the control side compared with the posterior group (P < 0.0001, Mann-Whitney). There was no significant difference between the control sides of the anterior and posterior groups (P = 0.223, Mann-Whitney). To make sure the differences we observed between the anterior and posterior groups were not related to variable time since onset, we normalized the GCC thickness by time (months since onset) and found that in anterior group, the rate of GCC thinning was much faster compared with the posterior group (anterior: 4.6 ± 1.1 µm/month, N = 13; posterior: 0.2 ± 0.5 µm/month, N = 36; P = 0.0002, Mann-Whitney) (Fig 2C). In 7 patients with congenital or incidentally noted occipital atrophy, there was 20.0 µm greater thinning in the abnormal side compared with the control side (abnormal: 58.1 ± 4.3 µm, control: 78.1 ± 3.0 µm, N = 7, P = 0.0156, Wilcoxon), which was not as severe as the anterior group but more severe than the acquired posterior group.

**Fig 2:**
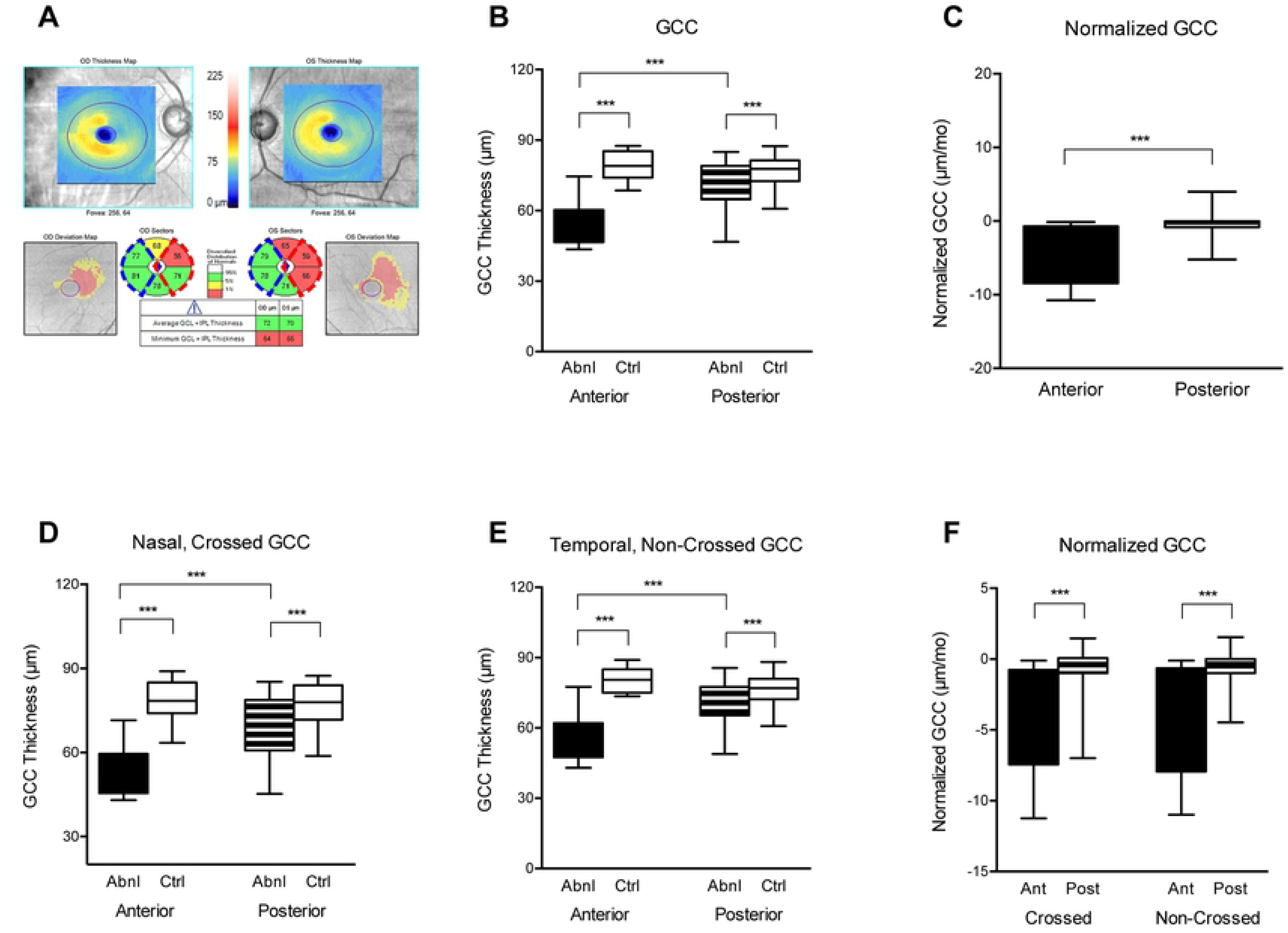
Significantly greater GCC thinning corresponding to visual field loss in the anterior group compared with the posterior group. **A.** An example of OCT ganglion cell analysis. The averaged red wedges represent “abnormal” and the averaged blue wedges represent “control” GCC thickness in a patient with right homonymous visual field defect. **B-F.** Graphs showing GCC thickness and rate of GCC thinning (OU, crossed, and non-crossed) for patients with anterior versus posterior visual pathway lesions. All graphs show significant differences for GCC between anterior and posterior groups. Error bars represent standard error of mean. *P between 0.01 to 0.05, **P between 0.001 to 0.01, and ***P < 0.001.

We asked whether there was a difference in retrograde degeneration of the crossed and the non-crossed pathways after acquired lesions of the visual pathway, since the crossed pathway is slightly longer [35, 36]. On macular GCC measurements, there was no significant difference in thinning when comparing the crossed vs. non-crossed pathway. There was 3.1 times as much crossed GCC thinning in the anterior group compared to the posterior group, and 3.5 times as much non-crossed GCC thinning in the anterior versus posterior group (Table 2) (Fig 2D-E).

**Table 2:** Amount of Crossed and Non-Crossed GCC and RNFL Thinning.

|  | Anterior |  |  |  | Posterior |  |  |  | P-value |
| --- | --- | --- | --- | --- | --- | --- | --- | --- | --- |
|  | N | Abnormal (μm) | Control (μm) | Dif (μm) | N | Abnormal (μm) | Control (μm) | Dif (μm) |  |
| **Crossed GCC** | 15 | 55.1 ± 2.6 | 79.0 ± 1.8 | 23.9 | 41 | 69.0 ± 1.7 | 76.7 ± 1.3 | 7.7 | **<0.0001<sup>a</sup>**<br>**<0.0001<sup>b</sup>**<br>**<0.0001<sup>c</sup>** |
| **Non-Crossed GCC** | 15 | 57.9 ± 2.6 | 80.3 ± 1.4 | 22.4 | 41 | 70.0 ± 1.5 | 76.4 ± 1.1 | 6.4 | **<0.0001<sup>a</sup>**<br>**<0.0001<sup>b</sup>**<br>**<0.0001<sup>c</sup>** |
| **Crossed RNFL** | 14 | 57.7 ± 2.4 | 69.4 ± 2.9 | 11.6 | 42 | 65.7 ± 1.7 | 66.4 ± 1.9 | 1.5 | **0.0284<sup>a</sup>**<br>**0.2810<sup>b</sup>**<br>**0.0078<sup>c</sup>** |
| **Non-Crossed RNFL (S/I)** | 14 | 87.5 ± 5.4 | 99.2 ± 4.0 | 11.7 | 42 | 105.2 ± 2.7 | 107.7 ± 2.5 | 1.6 | **0.0017<sup>a</sup>**<br>**0.3900<sup>b</sup>**<br>**0.0055<sup>c</sup>** |
| **Non-Crossed RNFL (S/I/T)** | 14 | 74.7 ± 4.6 | 83.9 ± 3.4 | 9.2 | 42 | 90.6 ± 2.1 | 92.7 ± 2.0 | 2.1 | **0.0012<sup>a</sup>**<br>**0.4800<sup>b</sup>**<br>**0.0047<sup>c</sup>** |

Dif: difference
<sup>a</sup> Wilcoxon between Anterior Abnormal and Anterior Control
<sup>b</sup> Wilcoxon between Posterior Abnormal and Posterior Control
<sup>c</sup> Mann-Whitney between Anterior Amount of Thinning and Posterior Amount of Thinning

**Table 3.** Rate of Thinning (µm/month) of Crossed and Non-Crossed GCC and RNFL.

|  | Anterior | Posterior | P-value<sup>a</sup> |
| --- | --- | --- | --- |
| **Crossed GCC** | 4.7 (N = 13) | 0.7 (N = 35) | **0.0004** |
| **Non-crossed GCC** | 4.5 (N = 13) | 0.6 (N = 35) | **0.0002** |
| **Crossed RNFL** | 2.3 (N = 12) | 0.6 (N = 34) | 0.1176 |
| **Non-Crossed RNFL**<br>**(S/I)** | 1.5 (N = 12) | 0.7 (N = 32) | 0.076 |
| **Non-Crossed RNFL**<br>**(S/I/T)** | 1.0 (N = 12) | 0.7 (N = 32) | 0.053 |

<sup>a</sup> Mann-Whitney between Anterior Rate of Thinning and Posterior Rate of Thinning

### Majority of GCC thinning occurred within 6 months

Given the difference in rate of thinning of GCC between the anterior and the posterior groups, we looked at GCC data only within 2 years of onset. The anterior group exhibited GCC thinning by the first OCT measurement, which was as early as 2 months after lesion onset. Within 4 months, the anterior group showed significant 19.9 µm of GCC thinning (abnormal: 60.5 ± 2.0 µm, control: 80.5 ± 2.2 µm, N = 6, P = 0.0313, Wilcoxon signed-rank test) (Table 4) (Fig 3A). The posterior group showed significant 3.7 µm of GCC thinning at 24 months after lesion onset (abnormal: 72.6 ± 2.0 µm, control: 76.3 ± 1.6 µm, N = 23, P = 0.0416, Wilcoxon signed-rank test). However, by 6 months after lesion onset, the majority of GCC thinning had already occurred in both groups (anterior: 78%, posterior: 58%).

**Fig 3:**
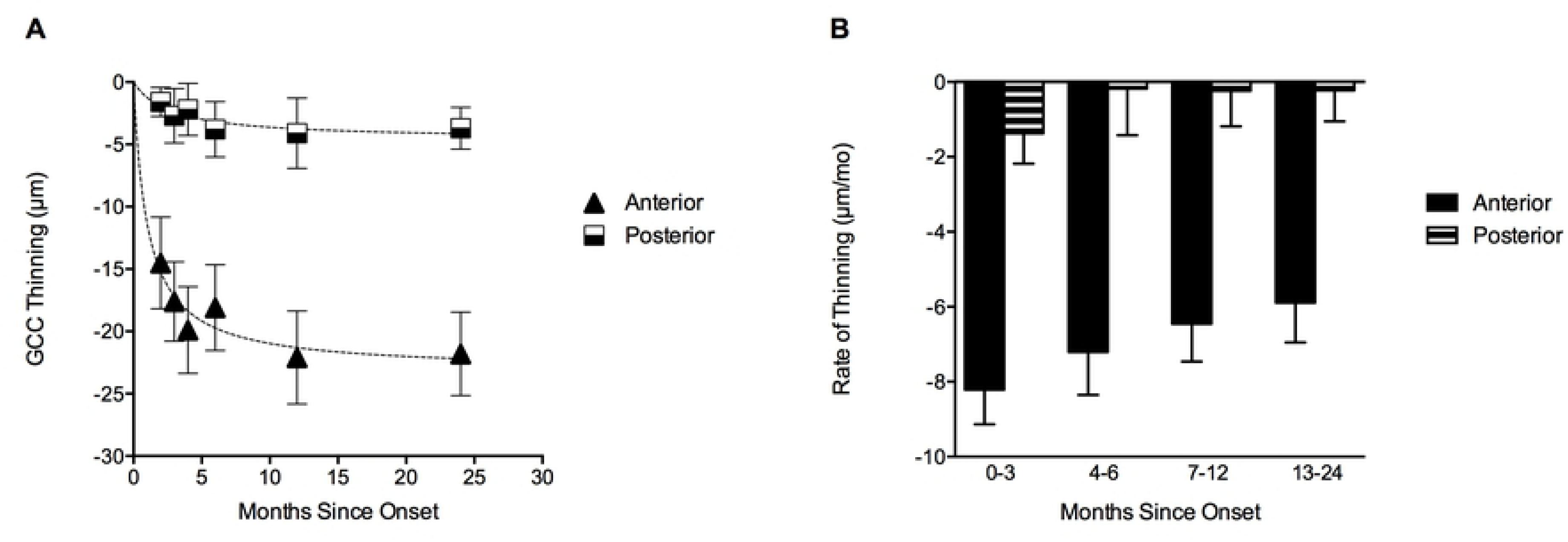
Timing and rate of macular GCC thinning in anterior and posterior groups. **A.** Cumulative difference between abnormal and normal GCC measurements in the anterior and posterior groups 2-24 months after onset. **B.** Bar graph showing rate of GCC thinning, calculated as cumulative difference between abnormal and normal GCC measurements normalized by time in months (mo). We binned the rate of thinning to show pattern of change at 0-3, 4-6, 7-12, and 13-24 months. All error bars represent standard error of mean.

**Table 4.** Time course of GCC thinning.

|  | Anterior |  |  |  | Posterior |  |  |  | P-value |
| --- | --- | --- | --- | --- | --- | --- | --- | --- | --- |
|  | N | Abnormal (μm) | Control (μm) | Dif (μm) | N | Abnormal (μm) | Control (μm) | Dif (μm) |  |
| < 2 mo | 3 | 63.3 ± 3.3 | 77.8 ± 2.4 | 14.5 | 9 | 73.6 ± 2.6 | 75.2 ± 2.8 | 1.6 | 0.2500<sup>a</sup><br>0.2344<sup>b</sup><br>**0.0182<sup>c</sup>** |
| <3 mo | 5 | 61.5 ± 2.1 | 79.1 ± 2.0 | 17.6 | 11 | 71.7 ± 3.4 | 74.5 ± 2.4 | 2.7 | 0.0625<sup>a</sup><br>0.3105<sup>b</sup><br>**0.0060<sup>c</sup>** |
| <4 mo | 6 | 60.5 ± 2.0 | 80.5 ± 2.2 | 19.9 | 12 | 72.3 ± 3.2 | 74.5 ± 2.2 | 2.2 | **0.0313<sup>a</sup>**<br>0.5625<sup>b</sup><br>**0.0014<sup>c</sup>** |
| < 6 mo | 7 | 62.1 ± 2.3 | 80.3 ± 1.9 | 18.1 | 15 | 71.8 ± 2.9 | 75.6 ± 2.0 | 3.8 | **0.0156<sup>a</sup>**<br>0.1879<sup>b</sup><br>**0.0023<sup>c</sup>** |
| < 12 mo | 9 | 59.6 ± 2.4 | 81.7 ± 1.7 | 22.1 | 20 | 72.2 ± 2.2 | 75.9 ± 1.7 | 4.1 | **0.0039<sup>a</sup>**<br>0.0531<sup>b</sup><br>**<0.0001<sup>c</sup>** |
| < 24 mo | 10 | 59.4 ± 2.2 | 81.2 ± 1.6 | 21.8 | 23 | 72.6 ± 2.0 | 76.3 ± 1.6 | 3.7 | **0.0020<sup>a</sup>**<br>0.0416<sup>b</sup><br>**<0.0001<sup>c</sup>** |
| All | 15 | 56.5 ± 2.6 | 79.6 ± 1.5 | 23.1 | 47 | 69.9 ± 1.6 | 76.5 ± 1.2 | 6.6 | **<0.0001<sup>a</sup>**<br>**<0.0001<sup>b</sup>**<br>**<0.0001<sup>c</sup>** |

Dif: difference
<sup>a</sup> Wilcoxon between Anterior Abnormal and Anterior Control
<sup>b</sup> Wilcoxon between Posterior Abnormal and Posterior Control
<sup>c</sup> Mann-Whitney between Anterior Amount of Thinning and Posterior Amount of Thinning

### Stable GCC thinning beyond 2 years except in patients with multiple insults

To determine whether more thinning occurred beyond 2 years, we examined 8 posterior lesion patients (total 10 events) who had 3 or more serial OCT measurements beyond 2 years (Fig 4B). Those with an isolated unilateral or bilateral occipital lobe event demonstrated little to no (≤ 2 µm) further thinning beyond 2 years of lesion onset (all dashed lines in Fig 4B). In comparison, 2 patients with metastatic cancer who underwent radiation therapy and more than one surgical resection were noted to have the largest amount of progressive macular thinning (7 µm and 18 µm decrease; solid orange and black lines in Fig 4B). One patient with an atypical venous stroke affecting the posterior temporal lobe had progressive thinning of 6 µm within 2-3 years, but no thinning beyond 3-4 years of onset (solid lime green line in Fig 4B). One patient with bilateral recurrent strokes showed progressive GCC thinning over time; there was 4 µm thinning within the first 3 years of a right occipital stroke and 8 µm thinning around 2-4 years after a left-sided occipital stroke (solid red and light blue lines in Fig 4B). Her brain MRI showed multiple areas of involvement consistent with multiple stroke events, which may account for the progressive GCC thinning seen over time. In the anterior group, 2 patients had progressive GCC thinning (3 µm and 6 µm decrease; solid red and maroon lines in Fig 4A). Both of the latter patients had multiple lesions over time.

**Fig 4:**
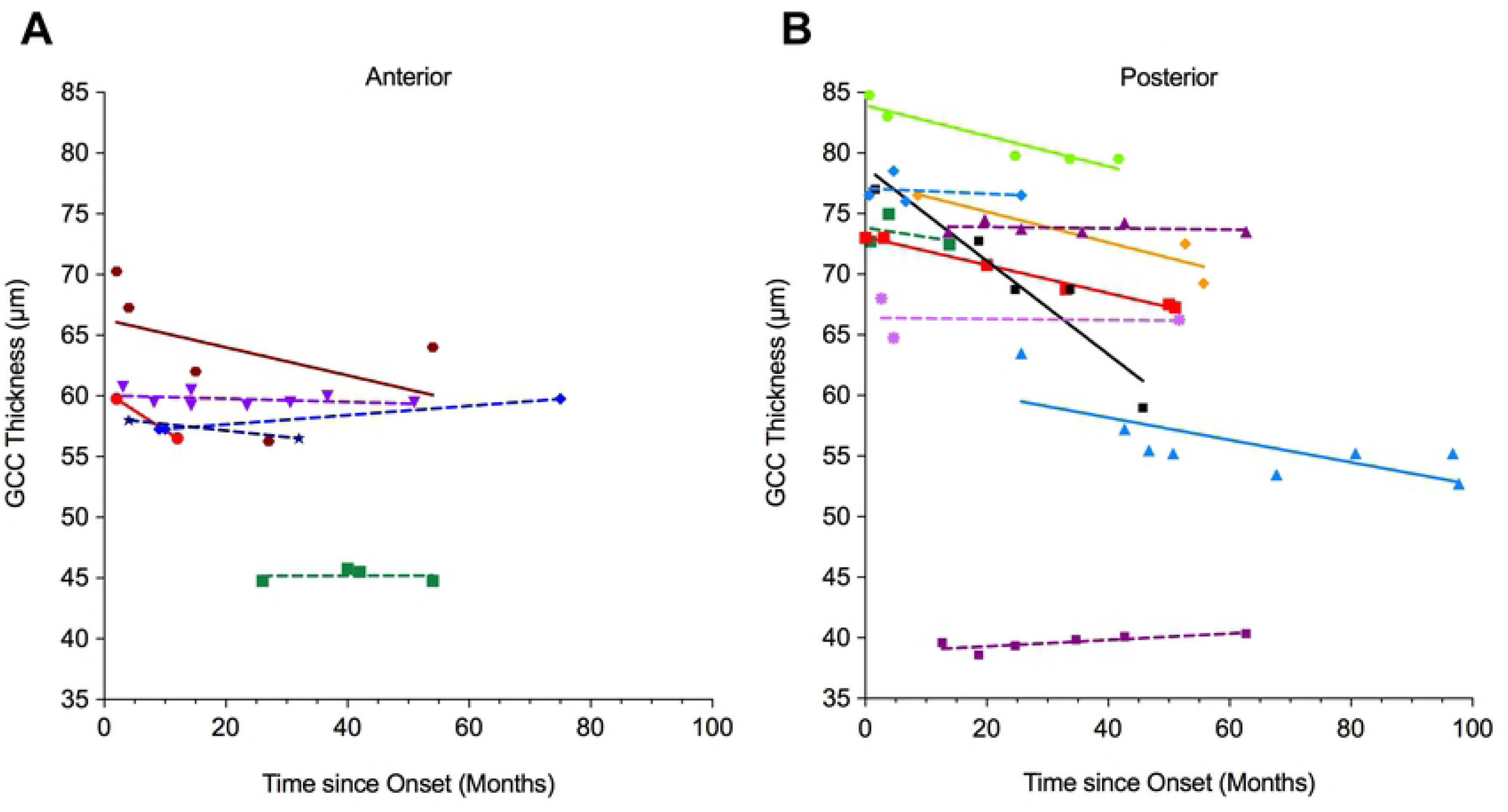
Serial GCC in anterior and posterior patients showed that many had stable GCC over years while some had further thinning. **A.** Macular GCC corresponding to the homonymous hemianopia in 6 anterior patients. Four had stable “abnormal” GCC thickness (dashed lines), while 2 had greater than 2 µm thinning (solid lines). **B.** Macular GCC corresponding to the visual field loss in 10 eyes in 8 patients in the posterior acquired group. Five events had stable GCC over years (dashed lines), while 5 events had progressive thinning beyond 2 years of lesion onset (solid lines; see details in Results).

### RNFL thinning showed similar pattern of thinning as GCC

To determine whether there is a difference in thinning in the unmyelinated axonal layer compared with the retinal layer containing the retinal ganglion cells, we first performed an analysis of the crossed RNFL (nasal quadrants) pathway; significant thinning was seen in both the anterior and posterior groups (Table 2). The anterior group had significantly 7.7 times greater crossed RNFL thinning compared with the posterior group (Fig 5B). In the non-crossed RNFL (superior and inferior quadrants) pathway, there was significantly greater non-crossed RNFL thinning in the anterior group compared with the posterior group (Table 2) (Fig 5D-5E). The amount and rate of RNFL thinning was similar to what was seen in the GCC analysis in both the anterior and posterior groups. To confirm this, we correlate GCC and RNFL measurements using a Gaussian nonlinear fit. We analyzed the normal and abnormal thickness of the crossed pathway (nasal GCC vs. nasal quadrant of RNFL) and found strong correlation between GCC and RNFL measurements (r^2^ = 0.953, N = 59, P = 0.985).

**Fig 5:**
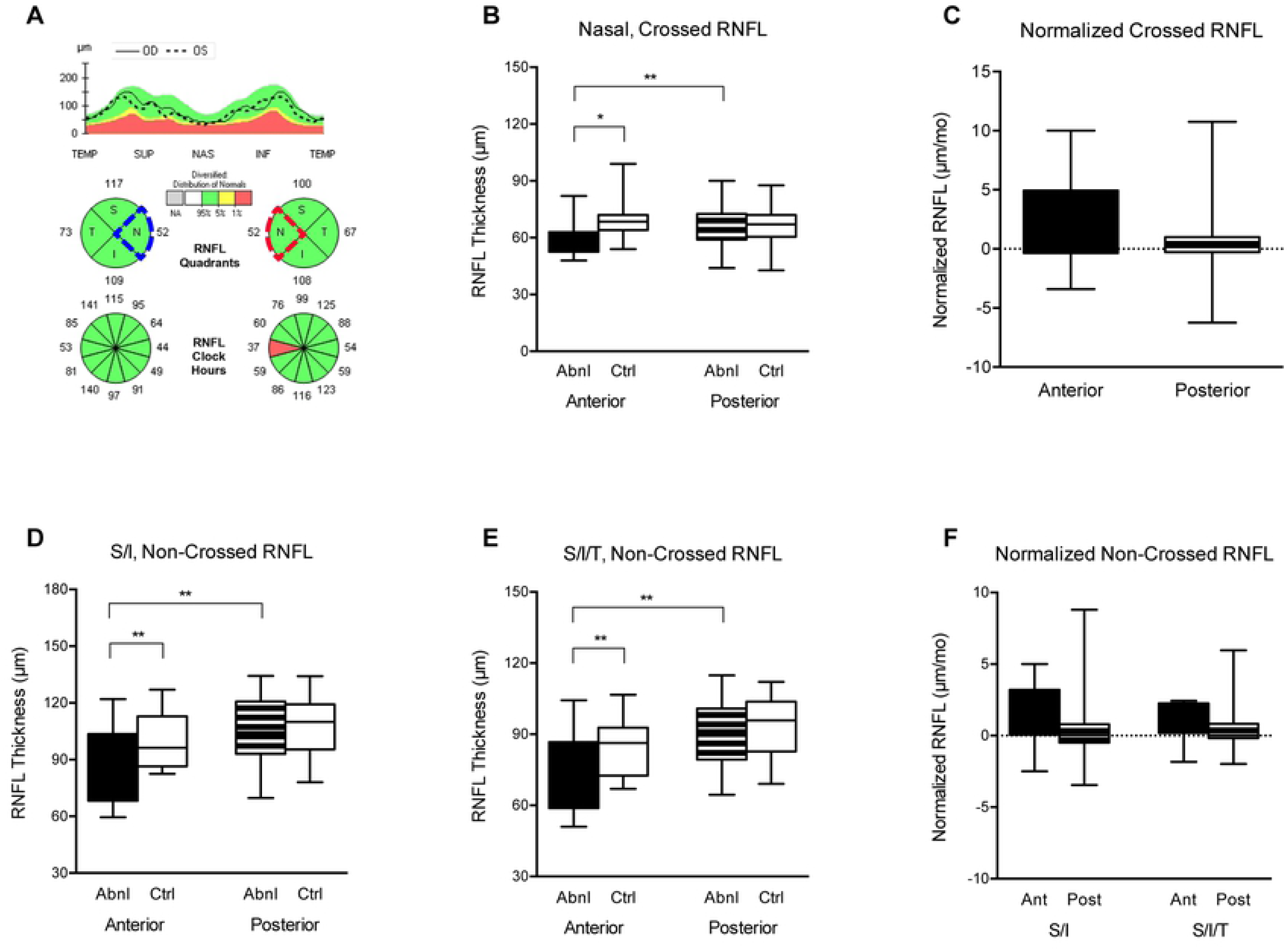
Greater RNFL thinning in anterior compared with the posterior group, and no difference in thinning between crossed and non-crossed pathways. **A.** Illustration of OCT retinal nerve fiber layer analysis. Blue wedge is the “control” and red wedge, the “abnormal” crossed RNFL quadrant in a patient with left homonymous visual field defect. **B.** Box-whisker plot of abnormal and control nasal crossed RNFL. **C.** Measurements in *B* normalized by time since onset in months (mo). **D.** Box-whisker plot of abnormal and control non-crossed RNFL, calculated as average of superior and inferior quadrants (S/I). **E.** Box-whisker plot of abnormal and control non-crossed RNFL, calculated as average of superior, inferior, and temporal quadrants (S/I/T). **F.** Measurements in *D* and *E* normalized by time since onset in months (mo). Error bars represent standard error of mean. *P value between 0.01 to 0.05, **P value between 0.001 to 0.01, and ***P value less than 0.01.

## Discussion

We performed a longitudinal study of anterior and posterior post-chiasmal lesions in patients with homonymous hemianopia to compare the severity and timing of retrograde degeneration of the retino-geniculo-cortical pathway. We found that anterior post-chiasmal visual pathway lesions led to prominent homonymous hemi-macular thinning within 2 months. One patient even had 9 µm thinning within one month. Within 3 months, the anterior group had 18 µm thinning or average rate of 8 µm per month. Within 6 months of onset, the anterior group exhibited about 5 times more GCC thinning than those with acquired lesions of the posterior visual pathway. *In contrast*, the posterior group had only 1.6 µm thinning at 2 months and 2.7 µm thinning at 3 months. At 3 months, posterior group had average rate of 1.4 µm per month, which was 33% that of the anterior group. After 6 months, there was relatively less thinning in both groups. Beyond 2 years, there was less than 2 µm thinning in both groups, except in a small number of patients who had multiple insults. Patients with congenital or incidentally noted posterior lesions had greater GCC thinning than those with acquired posterior lesions but less than patients with acquired anterior lesions. Both GCC and RNFL thinning were much more prominent and occurred earlier in the anterior group.

Although our study showed that both GCC and RNFL demonstrated similar patterns of degeneration, we found GCC analysis was better suited to study patients with homonymous hemianopia compared to RNFL analysis because of the clear segregation of neurons representing the nasal versus temporal visual fields. GCC analysis centers over the fovea and divides the crossed and non-crossed fibers into the nasal and temporal quadrants, making it simpler to classify abnormal and normal measurements. In our study, all GCC analysis (averaged, crossed, and non-crossed) was associated with more severe thinning and corresponded more strongly to homonymous visual field defect compared to all RNFL analysis (crossed, non-crossed). We observed that the averaged GCC, crossed GCC, and non-crossed GCC were similarly effective in examining retrograde degeneration. This is consistent with prior reports in the literature of patients with visual pathway lesions [33, 34, 37]. Comparison of timing of thinning of RNFL and GCC has also been studied in optic neuropathy patients, and GCC thinning has been reported earlier than RNFL thinning in disease processes like retrobulbar optic neuritis, papilledema, and anterior ischemic optic neuropathy [38, 39]. However, the GCC analysis is common to the Cirrus SD-OCT machine, which may not be consistently available in all eye clinics.

Following a lesion affecting the anterior post-chiasmal visual pathway, we found that retrograde non-trans-synaptic degeneration occurred significantly within 4 months and even as early as 2 months. Our findings are similar to what was found in previous small-scale human studies of anterior visual pathway lesions. Kanamori et al. studied 4 patients with optic tract lesion and noted mean 5.75 µm GCC thinning between 1 and 4 months after lesion onset, when comparing abnormal to control hemi-retinas [40]. A longitudinal study by Gabilondo et al. reported 27.5 µm of GCC thinning between abnormal and control hemi-retinas 5 months after optic tract lesion in a multiple sclerosis patient [41]. In our study, we showed that after damage to the post-chiasmal anterior visual pathway, which included the optic tract, mean 14.5 µm of GCC thinning occurred within 2 months. Because there is only one neuronal synapse in between the lateral geniculate nucleus and retinal ganglion cells, lesions of the anterior visual pathway directly affect the retinal ganglion cell axons. Meanwhile, between the striate cortex and RGCs lies the lateral geniculate-striate synapse, so lesions of the posterior visual pathway likely receive neurotrophic support from neuronal connections at both ends. This theory of neurotrophic support is currently our best explanation for why lesions of the post-chiasmal anterior visual pathway are associated with more severe and rapid retinal thinning compared to lesions of the posterior visual pathway. We propose that the presence of severe hemi-macular degeneration on OCT may be useful to clinicians in localizing lesions to the post-chiasmal anterior visual pathway.

Evidence of retrograde trans-synaptic degeneration after posterior visual pathway damage has been historically controversial because of difficulties in performing large pathologic studies in humans. In contrast, there is clear evidence in a small number of animal studies that retinal thinning after occipital lesion can occur substantially as early as 1 year. A study of 2 adult New World marmoset monkeys by Hendrickson et al. reported 20% ganglion cell loss based on photomicrograph analysis of retinal section at 1 year after visual cortex lesion [15]. Cowey et al. studied 17 macaque monkeys and found that the RGC count ratio between abnormal to control hemiretina (based on histologic section of the eye contralateral to brain lesion) was 0.7 at 1 year, 0.4 at 2.5 years, and 0.4 at 8 years after unilateral striate cortex ablation [42]. Interestingly, though our study was performed *in vivo* in humans, the timing of retinal thinning after visual cortex lesion is quite similar between our study and the non-human primate studies [15, 42]. Furthermore, we reported that on longitudinal follow-up, patients with occipital lobe damage did not demonstrate further retinal thinning after 2 years, which is consistent with the report by Cowey et al. of little to no RGC loss after 2.5 years after striate cortex ablation [42].

Human *in vivo* studies that used RNFL monitor retrograde trans-synaptic degeneration after occipital insults had variability in the severity of thinning, likely because of differences in methodology. We reported 3.1 µm of crossed RNFL thinning within 2 years, which we calculated by comparing abnormal and control hemiretinas between each individual patient (contralateral nasal retina for abnormal, ipsilateral nasal retina for control). Jindahra et al. studied 26 posterior lesion patients and reported average RNFL measurements of each eye; they observed 21.2 µm RNFL thinning in the crossed eye and 18 µm in the non-crossed eye, when compared to eyes from control patients [43]. Park et al. studied 46 patients with cerebral infarction affecting either the posterior cerebral artery, middle cerebral artery, or anterior cerebral artery, and reported 17.3 µm of crossed RNFL thinning [44] (when calculated using our paper’s methodology). Park et al. noted that time after stroke onset was significantly associated with reduced mean RNFL thickness, but did not further describe timing of when retrograde trans-synaptic degeneration first occurred. The majority of patients in the Park study (30 patients, 65%) had RNFL measurements taken after 2 years and up to 20 years from lesion onset. Perhaps the greater severity of crossed RNFL thinning reported by Park et al. is because of RNFL being measured much longer after lesion onset when compared to our study. It could also be due to their inclusion of patients with lesion of not only posterior cerebral artery territory but also middle and anterior cerebral artery territories. Gunes et al. studied 45 patients after MCA or PCA stroke and found little to no crossed RNFL thinning (-1.9 µm) [45], and Herro et al. noted 0 µm of crossed RNFL thinning in 9 occipital stroke patients [33]. The RNFL measurements from the latter two studies [33, 45] were calculated by the same methodology as our study and are more consistent with the changes our study found.

Human studies of patients with occipital lobe lesion have reported GCC measurements comparable to our finding of 4.1 µm of GCC thinning at 1 year of lesion onset. Anjos et al. studied 12 posterior cerebral artery stroke patients and found 15.5 µm of GCC thinning at 4.3 years after lesion onset when comparing abnormal to control hemiretinas of each patient (methodology identical to our study) [46]. A study by Herro et al. of 9 patients between 3-14 months after occipital stroke reported 5.3 µm of GCC thinning between affected and unaffected hemi-retinas [33]. In general, as we discussed earlier, the decreased severity of RNFL thinning in comparison to severity of GCC thinning provides further evidence that GCC is superior to RNFL for studying retrograde degeneration of the visual pathway [33, 37].

A review by Dinkin et al. suggests that understanding the time course of retrograde trans-synaptic degeneration could help clinicians determine the time of onset of brain lesions found incidentally on imaging and prognosticate visual field loss over time in patients with visual pathway damage [47]. Our study followed patients longitudinally by tracking OCT measurements over time with some patients being followed for more than 5 years, much longer than previously published studies on retrograde degeneration of the visual pathway [48]. We found 7.2 µm/year of nasal quadrant crossed RNFL thinning in posterior patients, which was comparable to Jindahra’s report of 4.4 µm/year of average RNFL thinning of the crossed *eye* [48]. We found there was a steady worsening of GCC thinning from 2, 4, 6, to 12 months (1.6 µm, 2.2 µm, 3.8 µm, and 4.1 µm of thinning, respectively) after onset of posterior visual pathway lesion. After 2 years, however, most of these patients did not exhibit further increase in severity of GCC thinning over time. The patients that did continue to demonstrate increasingly severe thinning beyond 2 years were the ones with multiple insults to the visual pathway. This progressive trend of GCC thinning is especially concerning and clinically important in patients with metastatic cancer, multiple neurologic surgeries, radiation treatments, recurrent stroke, and other insults. The progressive nature of retrograde degeneration in these patients with multiple insults are occurring at a measurable scale and may be occurring at a smaller scale in other patient groups with multiple insults, such as in patients with multiple concussions or multiple sclerosis.

A limitation of this study is that it was conducted retrospectively, and longitudinal measurements were limited by the number of OCT scans each patient already had in their medical records. However, this meant we could include a relatively large number of patients compared with previous publications, which provides a more comprehensive view of patients with homonymous hemianopia. The study was also limited because patients had diverse clinical presentations and severity of visual field loss. However, this broader approach was also done in historically important, large studies of homonymous hemianopia [1–4]. Just as the study by Zhang et al. [4] was superior because they used brain MRI to confirm lesion and provide anatomic correlation, our study provides an additional dimension of measurement by adding OCT and long-term follow-up to further understand this common and debilitating cause of vision loss.

## Conclusions

Our study fills an important gap in our understanding of long-distance retrograde degeneration by comparing the severity, timing, and rate of GCC and RNFL thinning in a large number of patients with homonymous visual field defect. A relatively large human study of retrograde degeneration is only possible due to advances in OCT imaging and its ability to provide high-resolution, reproducible, quantitation of the visual pathway at point-of-care locations. We found that anterior post-chiasmal lesions led to severe thinning of the retino-geniculate pathway within 2 months, with about 5 times greater thinning and 3 times faster rate of thinning at 3 months. We confirmed that trans-synaptic degeneration does occur after insult to the occipital lobe. Our study may be relevant to what also occurs following insults to other long white matter pathways. Finally, our study provides a global understanding of long-distance white matter tract degeneration, which is particularly important in consideration of the timing and type of therapies targeting regeneration or functional recovery in patients with homonymous hemianopia [49, 50].

## Acknowledgments

The authors have no acknowledgments.

